# A fast and cost-effective microsampling protocol incorporating reduced animal usage for time-series transcriptomics in rodent malaria parasites

**DOI:** 10.1101/350546

**Authors:** Abhinay Ramaprasad, Amit Kumar Subudhi, Richard Culleton, Arnab Pain

## Abstract

The transcriptional regulation occurring in malaria parasites during the clinically important life stages within host erythrocytes can be studied *in vivo* with rodent malaria parasites propagated in mice. Time-series transcriptome profiling commonly involves the euthanasia of groups of mice at specific time points followed by the extraction of parasite RNA from whole blood samples. Current methodologies for parasite RNA extraction involve several steps and when multiple time points are profiled, these protocols are laborious, time consuming, and require the euthanisation of large cohorts of mice. We designed a simplified protocol for parasite RNA extraction from blood volumes as low as 20 microliters (microsamples), serially bled from mice *via* tail snips and directly lysed with TRIzol reagent. Gene expression data derived from microsampling using RNA-seq were closely matched to those derived from larger volumes of leucocyte-depleted and saponin-treated blood obtained from euthanized mice and also tightly correlated between biological replicates. Transcriptome profiling of microsamples taken at different time points during the intra-erythrocytic developmental cycle of the rodent malaria parasite *Plasmodium vinckei* revealed the transcriptional cascade commonly observed in malaria parasites. Microsampling is a quick, robust and cost-efficient approach to sample collection for *in vivo* time-series transcriptomic studies in rodent malaria parasites.

## Background

High-throughput gene expression analysis is a powerful tool for profiling the transcription of thousands of genes at a particular point in time. Variations in gene expression in the malaria parasite across different life stages or conditions can reveal important aspects of gene regulation and function.

Transcriptome analyses of different life stages of *Plasmodium falciparum* have revealed that there is extensive transcriptional^1-4^ and post-transcriptional^5-8^ regulation of the genome^9^. Messenger RNA levels of various genes are observed to peak at different life stages during the intraerythrocytic developmental cycle (IDC) of the parasite, forming a patent transcriptional cascade in numerous *P. falciparum* strains^10^ and in other human malaria parasite species^11,12^. Such time-series transcriptome studies, including perturbation experiments^13-15^ can be performed with human malaria parasites but only in *in vitro* or *ex vivo* cultures. A few studies have profiled gene expression *in vivo* in clinical field isolates^16-18^ to infer gene function but gene expression changes due to a particular environmental condition or gene knockout need to be studied under controlled experimental settings.

Rodent malaria parasites (RMPs) allow for tractable *in vivo* model systems for the study of the biology of malaria parasites^19-21^. RMPs can be propagated in mice and mosquitoes under laboratory conditions, thus providing easy access to all the developmental stages of the parasite’s complex life cycle. Stage-specific transcriptional control has been shown in RMPs during their IDC^22-24^, vector^22,25-27^ and liver stages^28^. Thus, genome-wide transcription profiling in RMP models, in conjunction with manipulation of genetic or environmental factors of the host and/or the parasite, can provide valuable mechanistic insights into various aspects of parasite biology including antigenic variation and immunopathology^29-33^, vector transmission^34-37^ and drug resistance^38^.

Extraction of parasite RNA from blood stages of RMPs involves several steps. Peripheral, parasitized whole blood from infected mice is collected at a desired time point during the course of infection, through terminal sampling methods^39^ involving exsanguination. In the case of profiling life-stage specific gene expression in RMPs that exhibit asynchronous parasite development in the blood (*Plasmodium berghei* and *P. yoelii*), the parasite stages are typically separated via a density gradient column by centrifugation after blood collection or are maintained as synchronous *ex vivo* cultures^23, 24^

In order to study stage-specific gene expression in synchronous parasites such as *P. chabaudi* and *P. vinckei*, and more importantly, gene expression changes *in vivo* during the course of infection or in response to perturbations, transcriptomic profiling can be performed directly after blood collection. Blood samples are first enriched for parasite RNA by removing host leukocytes using either commercially available Plasmodipur filters (EuroProxima CAT#8011) or custom-made CF11 cellulose columns^40^. In addition to leukocyte depletion, host globin RNA within the parasitized erythrocytes is also removed by releasing the parasites from the RBCs via saponin lysis prior to RNA isolation. RNA is extracted from the purified parasites using guanidinium thiocyanatephenol-chloroform method^41,42^ (with TRIzol Reagent) or with column-based RNA extraction kits. RNA transcripts are then identified and their levels measured through microarray hybridization^22^ or next-generation sequencing of RNA-seq libraries^23, 24^

While profiles from both methods correlate well^3,43^, RNA-seq involves direct, deep sequencing of RNA transcripts and offers an unbiased and highly resolved picture of the parasite’s transcriptional landscape by providing information on genes expressed at low levels, alternative splicing events and antisense transcription at base-pair resolution^3,4,43,44^.

As these experiments require exsanguination of mice, large cohorts of mice are required for each time point (see Figure 1A). Thus, the number of animals that need to be euthanized directly increases with the number of biological replicates and time points, imposing ethical and cost constraints on the study design. Sampling different animals at different time points also increases the probability that inter-animal variation will increase the variability of the results.

**Figure-1.**
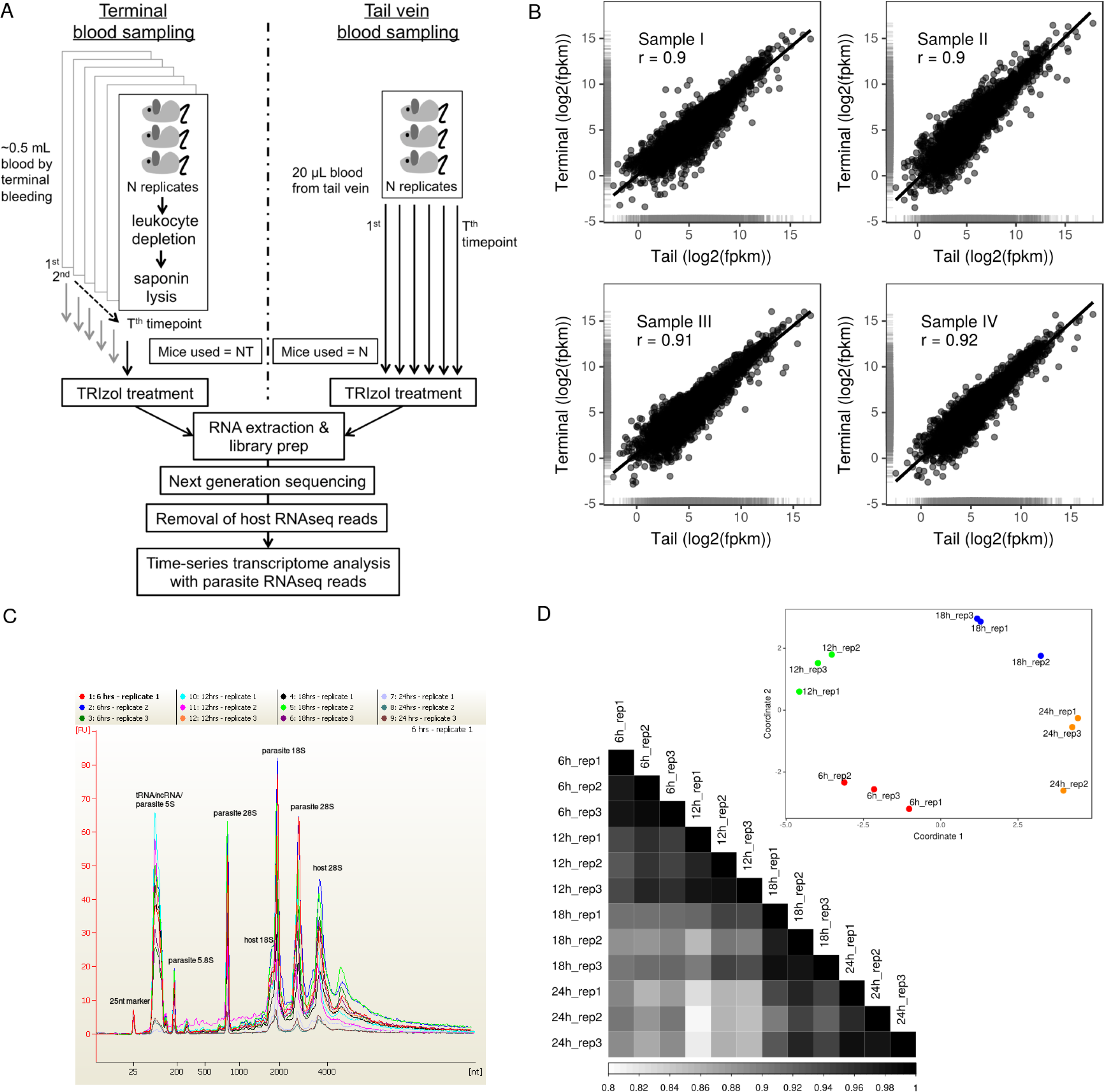
Microsampling protocol design and reproducibility. A) In terminal blood sampling, at each time point, groups of mice are exsanguinated to get 0.5 - 0.6 mL blood volumes, which is then subject to leukocyte depletion and saponin lysis before TRIzol treatment. Thus, the number of mice increases proportionally to number of time points and biological replicates in the study design (NT). On the other hand, microsampling involves obtaining sample volumes as low as 20 μL from the same mouse at different time points, thus confining the number of mice to just biological replicates (N) and significantly lowering costs and biological variability due to individual animals. Leukocyte depletion and saponin lysis are also not performed on the low volume samples, thus saving time and manpower. B) High Pearson correlations were observed between gene expression profiles from microsampling and terminal blood sampling methods. C) Bioanalyser electrophoregrams of total RNA from *P. vinckei* microsamples show that high quality RNA could be extracted consistently from 20 μL microsamples. D) Heatmap shows pair-wise Pearson correlation coefficients and the inset shows multidimensional scaling to visualize the level of similarity between the *P. vinckei* microsamples. Microsamples show low degree of variability and are highly reproducible as proved by tight correlations between biological replicates.

Exsanguination involves deep terminal anesthesia of the mouse, and the performance of surgical procedures. This, along with the leukocyte depletion and saponin lysis steps, makes the entire procedure time-consuming and requires considerable technical expertise. Thus, multiple sampling at short time intervals requires significant cost, time-investment and high level of technical expertise. We have, therefore, devised a simplified protocol for time-series transcriptomics of RMPs that uses a serial blood microsampling approach for sample collection (see Figure 1A).

Microsamples are usually blood volumes less than 50 μL and which can be collected at multiple time points from a single mouse using less invasive procedures such as tail snip or tail vein sampling. Microsampling techniques are quicker, cause less stress to the animal, allow multiple samples from the same animal through time and have been shown to reduce animal usage in pharmacokinetic studies^45-48^. Here, we have evaluated the feasibility of sequencing parasite RNA transcripts from blood volumes as low as 20 μL and assessed whether data thus obtained would reflect the true global gene expression in the parasite. We have also assessed the impact of raw processing of blood samples without leukocyte depletion.

## Results

Two sets of experiments were designed to assess the transcriptomic read-outs from the microsampling approach. First, we compared the gene expression profile obtained through microsampling with that of the routinely applied technique of terminal bleeding, in mice infected with *P. chabaudi* parasites. Twenty microlitres of blood was collected via tail snip from *P. chabaudi*-infected mice, washed with PBS and immediately lysed with 0.5 mL TRIzol reagent. Following microsample collection, mice were anaesthetized and exsanguinated *via* incision of the brachial artery. Around 0.5 mL of blood was collected from each mouse, washed twice with PBS and passed through home-made CF11 cellulose columns to remove leukocytes. The RBCs were then gently lysed using saponin and the harvested parasite pellet was immediately lysed with 1 mL TRIzol reagent.

In the second experiment, microsampling was done from mice infected with *P. vinckei* parasites to assess whether the microsampling protocol can identify gene expression changes that occur during *P. vinckei*’s intraerythrocytic developmental cycle. Twenty microlitres of samples (microsamples) were taken *via* tail snip from three mice infected with *P. vinckei* at four time points - 6h, 12h, 18h and 24h during the 4th day post infection, and lysed with TRIzol, as before.

Firstly, RNA yield and quality obtained from microsamples were assessed. The *P. chabaudi* samples were taken at a low parasitaemia of 4-8% and the *P. vinckei* samples at a higher parasitaemia of around 25%. This was reflected in the total RNA yield from these samples, with most of the *P. chabaudi* and *P. vinckei* samples yielding around 1 μg and 10 μg respectively (see Table 1). Quality assessment of the total RNA using Bioanalyser found no evidence of degradation and also, as expected, showed rRNA electropherogram peaks of both mouse and *Plasmodium* origin (see Figure 1C).

**Table 1.** Microsample characteristics and RNA-seq mapping statistics. Microsamples from *P. chabaudi* had low parasitaemia and therefore, a low percentage of reads mapping to *P. chabaudi* genome. In contrast, *P. vinckei* microsamples had lesser host contamination resulting in higher median fragment coverage across its transcripts. Transcript integrity number (TIN) was calculated using RSeQC ^56^ and all samples showed a high TIN value, indicating little to no evidence of RNA degradation.

| RMP | Microsample | Parasitaemia | RNA yield (µg) | Total Reads | Parasite reads | Mapping percentage | Median fragment coverage | Median TIN |
| --- | --- | --- | --- | --- | --- | --- | --- | --- |
| *P. chabaudi* | Sample I | 7.74 | 0.87 | 26,585,563 | 1,228,998 | 4.62 | 8 | 87.66 |
|  | Sample II | 6.22 | 0.42 | 19,842,987 | 2,458,755 | 12.39 | 49 | 86.41 |
|  | Sample III | 5.75 | 1.71 | 15,332,092 | 676,488 | 4.41 | 10 | 92.2 |
|  | Sample IV | 3.64 | 1.03 | 31,385,250 | 1,420,373 | 4.52 | 26 | 91.13 |
| *P. vinckei* | 6h_rep1 | 23.88 | 9.6 | 32,755,140 | 22,915,171 | 69.96 | 499 | 91.27 |
|  | 6h_rep2 | 22.19 | 9 | 31,427,694 | 21,274,632 | 67.69 | 479 | 91.15 |
|  | 6h_rep3 | 23.14 | 9.03 | 31,331,956 | 20,044,950 | 63.98 | 424 | 91.04 |
|  | 12h_rep1 | 24.19 | 11.49 | 35,255,928 | 26,290,657 | 74.57 | 710.5 | 91.79 |
|  | 12h_rep2 | 24.32 | 11.46 | 28,504,674 | 16,567,841 | 58.12 | 451.5 | 86.61 |
|  | 12h_rep3 | 22.75 | 10.14 | 27,368,020 | 17,521,494 | 64.02 | 483.5 | 92 |
|  | 18h_rep1 | 24.77 | 6.33 | 27,622,130 | 18,733,720 | 67.82 | 582 | 89.88 |
|  | 18h_rep2 | 25.65 | 11.49 | 29,253,240 | 18,590,556 | 63.55 | 552 | 91.11 |
|  | 18h_rep3 | 23.11 | 10.23 | 27,377,938 | 15,234,884 | 55.65 | 468 | 90.08 |
|  | 24h_rep1 | 27.06 | 5.4 | 33,624,482 | 25,522,094 | 75.90 | 698 | 91.57 |
|  | 24h_rep2 | 25.92 | 1.24 | 35,255,928 | 20,282,255 | 57.53 | 501.5 | 87.43 |
|  | 24h_rep3 | 26.45 | 1.95 | 35,255,928 | 23,045,897 | 65.37 | 634 | 90.69 |

Upon performing Illumina RNA sequencing and mapping RNA-seq reads onto the relevant RMP reference genomes, it was found that a low percentage of the reads (4-12%) were of parasite origin in the *P. chabaudi* samples, resulting in low fragment depth (most genes had less than 50 fragments mapped onto them) while higher percentages (55-75%) and fragment depth (most genes with more than 424 mapped fragments) were obtained for *P. vinckei* (see Table 1).

### Comparison of microsampling and terminal sampling methods

There was strong correlation between the normalized gene-wise FPKM values (fragments per kilobase of transcript per million mapped reads) of microsamples and terminal bleed samples (see Figure 1B). Pearson correlations of 0.9 to 0.92 were obtained in the four mice that were profiled, based on the expression values of 3,936 genes. Similar correlation was obtained independently by real-time quantitative PCR (qPCR) of 91 genes (see Supplementary Figure S2). However, the microsamples cumulatively failed to measure FPKM values for 942 genes compared to the terminal bleed samples, owing to their lower fragment depths (terminal bleed samples had a fragment depth higher than 100).

### Time-series transcriptomics in P. vinckei

*P. v. vinckei* CY is a synchronous parasite^49,50^ and the four time points sampled correspond to roughly dominant populations of ring (6h), early trophozoite (12h), late trophozoite (18h) and schizont (24h) stages. Multidimensional scaling of the samples from the four time points showed good level of dissimilarity between their expression profiles reflecting stage-specific gene expression in the parasite. Tight correlations were also obtained among the three biological replicates at all four time points (Pearson correlations ranging from 0.97 to 0.99) (see Figure 1D). Of a total of 5,073 genes in *P. vinckei*, 4,328 genes were significantly differentially expressed (p-value less than 0.05) in at least one time point (616 genes were not differentially expressed and only 129 genes had 0 FPKM value in at least one timepoint). As in other *Plasmodium* species^1,3,10,11,23,24^ stage-specific gene expression was inferred in *P. vinckei* by constructing a phaseogram, where the differentially expressed genes are ordered according to the time point at which their expression peaks (see Figure 2).

**Figure-2.**
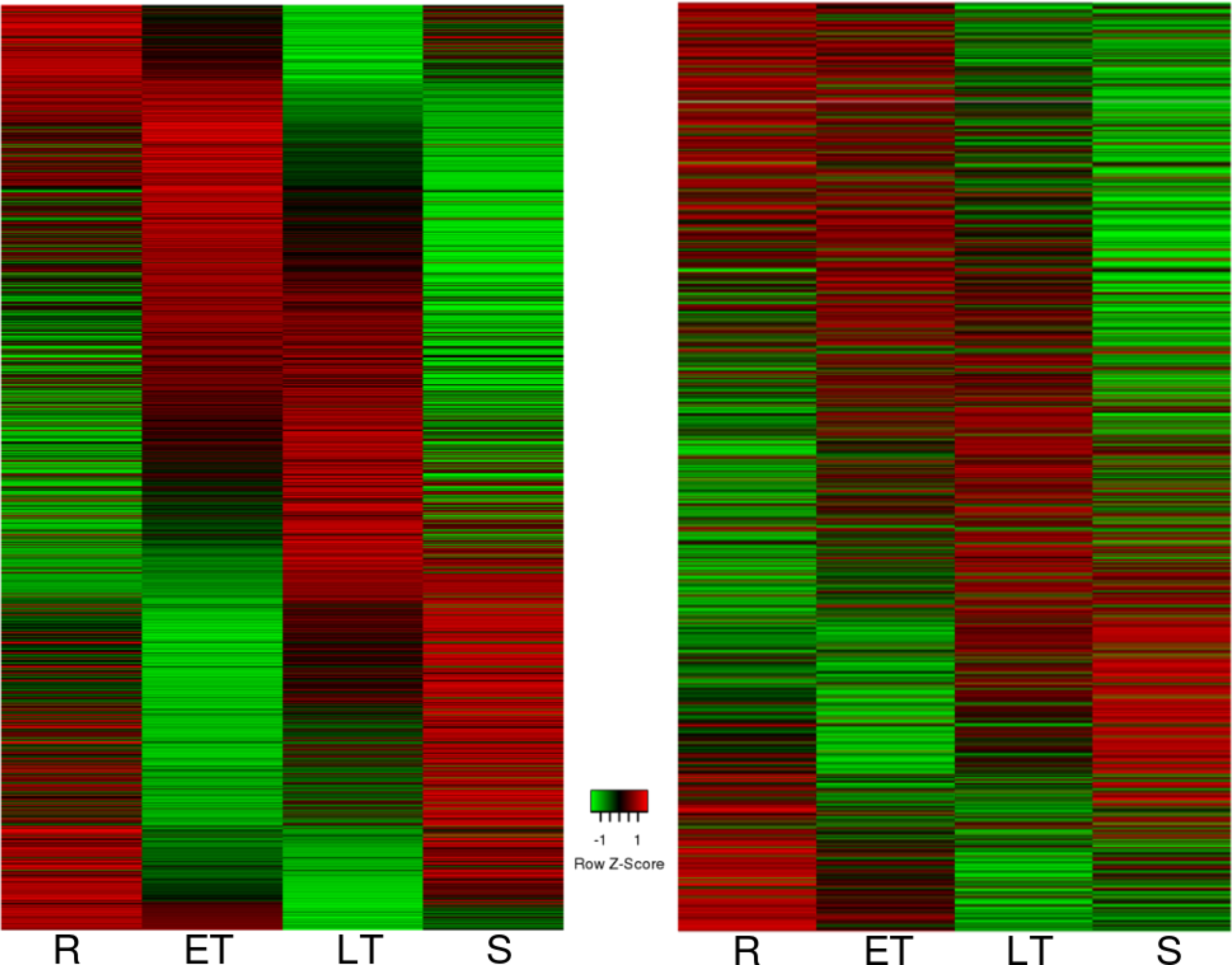
Time-series transcriptome of *P. vinckei vinckei* CY. Heat maps showing gene expression in *P. vinckei* at 6 hour time points during the 24 hour asexual cycle, each corresponding to a dominant population of rings (R), early trophozoites (ER), late trophozoites (LT) and schizonts (S) respectively. All significantly regulated *P. vinckei* genes (4,328 genes) were ordered according to their phase of expression (left). *P. vinckei* genes with one-to-one orthologs in *P. falciparum* (2,480 genes) were ordered based on *P. falciparum* gene expression pattern shown in ^1^ (right). Gene-wise FPKM values can be found in Supplementary tables S1 and S2.

Next, we compared *P. vinckei* gene expression with the transcriptional cascade shown in *P. falciparum* ^51^. Two thousand, four hundred and eighty (2,480) *P. vinckei* genes were ordered according to the expression values of their one-to-one orthologues in *P. falciparum* (from a total of 2,712 *P. falciparum* genes profiled in ^51^), and a similar temporal expression cascade as in 51 was obtained (see Figure 2).

### Minimum sequencing depth and cost estimation

The absence of any host depletion step reduces the proportion of sequencing reads of parasite origin and the final read coverage of parasite transcripts. In order to assess the impact of host contamination, we first estimated the minimum amount of sequencing required per sample to gain robust results during gene expression analysis.

Random subsampling of different sizes (1, 3.16, 10, 31.6 and 100% of the total reads) was performed for the 12 *P. vinckei* samples and differentially expressed genes were inferred in each case at a significance level (q-value) of 0.05. It was observed that the number of differentially expressed genes and their expression values (in all pairwise comparisons between the four time points) did not change drastically in subsamples of and above 31.6% of the total reads, which is equivalent to around 3 million paired-end reads per sample (see Supplementary Figure S1). Setting this as the target sequencing depth for an RMP transcriptome, sequencing and animal costs alone were calculated for different host contamination levels (10 to 80%) for a microsampling experiment and compared with a terminal blood sampling experiment. As the number of time points or biological replicates increase in the study design, microsamples with host contamination levels less than 70% would cost the same or less than terminal blood sampling (see Supplementary Figure S1). Differences in other costs incurred for animal housing and laboratory reagents between the two protocols were assumed to be negligible. Our estimates were also made without considering the substantial manpower costs associated with the terminal blood sampling procedure.

## Discussion

Serial profiling of gene expression during the course of infection of rodent malaria parasites can be a powerful tool for studying host-parasite interactions and gene regulation during the clinically important blood stages of the parasite. Blood sampling in time-series experiments are usually carried out through terminal techniques in order to obtain sufficient blood volumes for subsequent host leukocyte removal and for isolating large quantities of total RNA to satisfy the input requirements of microarray or sequencing protocols.

However, when the study design involves several time points or biological replicates, terminal blood sampling becomes laborious and requires large numbers of mice. Here we propose microsampling as a quick, easier, non-invasive alternative, which allows serial sampling of small blood volumes from the same animal. The microsampling of blood from mice has proved to provide robust results in pharmacokinetic studies ^45-48^, but its application for transcriptomic profiling has not been evaluated.

Given that only a small fraction of the parasite population is sampled during microsampling compared to terminal techniques, it is possible that the former method may provide a biased or highly variable gene expression profile. However, our experiments demonstrate that microsamples from biological replicates show highly similar expression profiles and also reflect closely the expression levels obtained from terminal blood sampling. We have shown correlations of 0.9-0.92 between microsamples and terminal bleed samples. It is possible that some of the variation observed may be due to the 20−30 minutes time lag between microsampling and terminal bleed points due to the anaesthetizing and exsanguination of mice for blood collection.

Gene expression analysis of microsamples collected from four time points or life stages across the^24^ hours life cycle of *P. vinckei* showed most of its genes differentially expressed and forming a transcriptional cascade typical of a malaria parasite. Moreover, orthologous genes between *P. vinckei* and *P. falciparum* showed similar expression profiles in their respective life stages. Thus, our protocol was able to capture the transcriptional regulation occurring in *P. vinckei* life stages but at lower cost, time and effort than previous protocols for profiling stage-specific gene expression in RMPs.

Our simplified approach offers several advantages over standard techniques. It drastically reduces the number of animals used. In the time-series experiment in *P. vinckei*, only three mice were used, where 12 mice (four time points and three biological replicates) would have been required in the case of terminal blood collection. This relaxes ethical and cost constraints on study designs. More time points and biological replicates could be included for performing transcriptome analysis and drawing conclusions with better statistical power. Microsampling is very quick and it takes less than five minutes to collect, wash and stabilize the sample in TRIzol. This reduces the time elapsed between sample collection and cell lysis, thus providing a “snapshot” of gene expression at a particular time point. While we have used 20 μL blood volumes here, with the availability of efficient RNA extraction and low-input RNA library preparation kits, it is also possible to process microsamples less than 20 μL.

Quick sampling and low sample volumes will enable gene expression profiling at more frequent time points. Microsampling techniques are less invasive and do not require warming of the animal, thus reducing animal stress. While we have used tail snip blood collection here, other suitable methods^52^ such as tail vein sampling, saphenous vein sampling and capillary microsampling^53^ can be adopted to further reduce animal stress. These collection methods are also simple and do not require expertise in surgical procedures.

Our protocol allows for expression profiling at multiple time points from the same host, thus reducing animal-to-animal variation.

As host RNA depletion steps can be skipped, high proportions of host-derived reads in the sequencing data is the main limitation of this protocol, especially at low levels of parasitaemia. More sequencing data is therefore required per sample to compensate for host contamination and to achieve a suitable sequencing depth of the parasite’s transcriptome. By randomly reducing the number of reads in our dataset, we estimated that only 3 million paired-end reads are required for robust differential expression analyses. Increasing the number of replicates could further reduce this minimum sequencing depth.

Around 20% of the genes did not have fragment coverage in microsamples with *P. chabaudi* at low parasitaemias suggesting that host contamination at parasitaemias of below 7% would be unfeasible, requiring extra-large amount of sequencing to achieve sufficient sequencing depth for parasite transcripts. The protocol is well-suited for higher parasitaemias as shown in the *P. vinckei* microsamples that yielded sufficient fragment coverage for almost all of the genes.

Based on current animal and sequencing costs, we estimate that any study with host contamination as high as 70% would still be economically viable, especially if the significant reduction in manpower costs is considered. Host reads would of course be informative in studies that profile both host and parasite transcriptomes simultaneously to study host response to infection.

In conclusion, RNA extraction, sequencing and expression analysis can be performed with <20 microliters of malaria parasite infected blood in a robust, reproducible and cost efficient way. Our protocol can also be adapted to profile the *in vivo* transcriptome of other blood-borne pathogens like Trypanosomes in rodent models. Blood collection and TRIzol lysis can be performed within 5 minutes allowing snapshots of gene expression to be taken quickly, at more frequent time points, and using less manpower. Serial bleeding of the same mice throughout the study reduces the number of animals used and animal-to-animal variation.

## Methods

### Laboratory animals and rodent malaria parasites

Six to eight weeks old female CBA mice (SLC Inc., Shizuoka, Japan) were used in all experiments. Mice were housed at 26°C and maintained on a diet of mouse feed (CLEA Rodent 499 Diet CE-2 from CLEA Japan, Inc.) and water. Mice infected with malaria parasites were given 0.05% para-aminobenzoic acid (PABA)- supplemented water to assist parasite growth.

Laboratory animal experimentation was performed in strict accordance with the Japanese Humane Treatment and Management of Animals Law (Law No. 105 dated 19 October 1973 modified on 2 June 2006), and the Regulation on Animal Experimentation at Nagasaki University, Japan. The protocol was approved by the Institutional Animal Research Committee of Nagasaki University (permit: 12072610052).

*Plasmodium chabaudi chabaudi* AS strain and *Plasmodium vinckei vinckei* CY strain were used to initiate infections in mice. In each case, one million parasites were intravenously inoculated into each CBA mouse.

### Blood sampling

#### Comparison of microsampling and terminal sampling methods

In order to compare microsampling with terminal bleed sampling, blood sampling was performed in mice infected with either wild-type *P. chabaudi* parasites or genetically modified *P. chabaudi* parasites (PCHAS_1433600 gene knockout). On the fourth day post infection, each mouse was restrained and 1-2 mm of the distal portion of the tail was excised with sanitized scissors. Twenty microlitres of blood was subsequently collected from the tail by pipette and deposited in 500 μL of phosphate buffered saline (PBS) solution. Whole blood was briefly spun down in a tabletop microcentrifuge, supernatant removed and the RBC pellet resuspended in 500 μL TRIzol reagent (ThermoFischer Cat#15596026). TRIzol lysates were temporarily stored at 4C (for periods up to 48hrs), or for longer periods at -80°C.

Thin blood smears on glass slides were taken just before blood collection of blood, fixed with methanol and stained with Giemsa’s solution for estimating parasitaemia.

Following this, mice were immediately anaesthetized with an intraperitoneal injection of 0.2 mL of 10% sodium pentobarbital solution in PBS solution. Once completely sedated, a vertical incision was made from the bottom of the rib cage to the right shoulder, forming a cavity. The brachial artery was cut and around 0.5-0.6 mL of blood was collected into 3 mL citrate saline solution (8.5g of NaCl,15g trisodium citrate in 1L of distilled water, pH 7.2) on ice with a sterile Pasteur pipette. The complete procedure was carried out with the mouse under isoflurane anaesthesia via inhalation. Mice were euthanized by cervical dislocation.

Parasitized blood was centrifuged at 2000 rpm for 5 min and the pellet washed once with 10 mL PBS solution to remove blood serum. The RBC pellet was obtained by further centrifugation at 2000 rpm for 5 min and was then resuspended in 10 mL PBS solution. Cellulose columns (Sigma Cat# C6288) were prepared and equilibrated with PBS solution, following which blood solution was passed through the column to deplete mouse leukocytes. RBCs were then gently lysed with 0.15% saponin solution, centrifuged at 3000 rpm for 5 min and ghost RBCs carefully removed leaving behind the parasite pellet. The parasite pellet was treated with 1 mL TRIzol reagent and stored at 4°C.

### Time-series transcriptomics

*Plasmodium vinckei* infections were initiated in three CBA mice. On day four post infection, 20 μL blood was collected *via* tail snip at four time points; 06:00 hrs, 12:00 hrs, 18:00 hrs and 24:00 hrs. Blood samples were processed as before and TRIzol lysates were stored at 4°C. Blood slides were taken just before blood collection and parasitaemia and proportions of different life stages (rings, early trophozoites, late trophozoites and schizonts) were measured. Mice were euthanized by cervical dislocation at the completion of the last sampling time point.

RNA extraction, library preparation and sequencing of RNA isolated from TRIzol was performed according to the manufacturer’s protocol (Invitrogen). The RNA pellet was resuspended in 15 μL nuclease-free water, RNA quantity measured by Qubit flourometer and RNA integrity measured by Agilent Bioanalyser (Agilent RNA 6000 Nano kit Cat#5067-1511).

Strand-specific mRNA sequencing was performed from total RNA using a TruSeq Stranded mRNA Sample Prep Kit LT (Illumina Cat#RS-122-2101) according to the manufacturer’s instructions. Briefly, polyA+ mRNA was purified from total RNA using oligo-dT dynabead selection. First strand cDNA was synthesized using randomly primed oligos followed by second strand synthesis where dUTPs were incorporated to achieve strand-specificity. The cDNA was adapter-ligated and the libraries amplified by PCR. Libraries were sequenced in an Illumina Hiseq2000 with paired-end 100 bp read chemistry.

### RNA-seq read mapping and gene expression analysis

Strand-specific RNA-seq paired-end reads were mapped onto the reference genomes, *P. chabaudi* AS version 3 (http://www.genedb.org/Homepage/Pchabaudi) and *P. vinckei vinckei* CY genome (ENA study accession - PRJEB27301) using TopHat2 v2.0.13 ^54^ with options “–library-type=fr-firststrand” and “–no-novel-juncs”. Differential expression analysis was carried out using cuffdiff2 v2.2.1 ^55^ with “-u -b” parameters. Transcript integrity number (TIN) was calculated from the mapped reads using RSeqC ^56^. Pearson correlation coefficients were calculated using *cor* function in R *stats* package ^57^ and visualized using *corrplot* package ^58^ in R. Multidimensional scaling was performed using *cmdscale* and *dist* functions in stats R package. To create a phaseogram, the phase of gene expression was calculated using the ARSER ^59^ package and the genes were ordered according to their phase. Heatmaps were created using *heatmap.2* function in *gplots* package ^60^ and graphs plotted using *ggplot2* ^61^ in R.

### Real-time qPCR using Biomark HD system

cDNA synthesis was performed using reverse transcription master mix according to the manufacturer’s instructions (Fluidigm). Pre-amplification of target cDNAs were performed using a multiplexed, target-specific amplification protocol (95°C for 15 sec, 60°C for 4 min for a total of 14 cycles). The pre-amplification step uses a cocktail of forward and reverse primers of genes of interest to increase the number of copies to a detectable level. Products were diluted 5 folds prior to amplification using SsoFast EvaGreen Supermix with low ROX and target specific primers in 96.96 Dynamic arrays on a Biomark HD microfluidic quantitative RT-PCR system (Fluidigm) (run as technical duplicates). Expression data for each gene was retrieved in the form of C_t_ values. The gene expression (in the form of dCt values using PCHAS_1202900 as housekeeping gene) of 91 genes were assessed and compared between 2-4 biological replicates of microsamples and terminally bled samples as a validation of comparisons done by RNA-seq.

### Subsampling and cost estimation

Gene-wise fragment counts were inferred using featureCounts ^62^ and subsampling analysis was performed using subSeq v1.4.1 ^63^. Random subsampling of different sizes (1, 3.16, 10, 31.6 and 100% of the total reads) was performed for the 12 *P. vinckei* samples with up to 10 replications at each subsampling step. For each subsampling step, differential expression analysis was performed for each pairwise comparison among the four time points with a q-value cutoff of 0.05 using two tools, DESeq2 ^64^ and edgeR ^65^. Cost estimation was done with the following conditions- i) target sequencing depth of 3,000,000 paired-end 100bp reads per sample, ii) sequencing cost per gigabasepair is $22 ^66^ and iii) Cost of one six-weeks old female CBA/J mouse is $31.68 (https://www.jax.org/strain/000656).

## Acknowledgements

The authors would like to acknowledge the Bioscience Core Laboratory (BCL) at King Abdullah University of Science and Technology for their help with next generation sequencing. RC was supported by a grant (JP16K21233) from the Japan Society for the Promotion of Science. This work was supported by KAUST baseline fund (BAS/1/1020- 01-01) and Competitive Research Fund (URF/1/2267-01-01) to AP.

## Author’s contributions

AR, RC and AP designed the methodology. AR collected, analyzed and interpreted the data. AKS designed and performed real-time qPCR experiments. AR wrote the manuscript and all authors contributed to it.

## Declarations

The authors declare that they have no competing interests. All raw sequencing fastq files are available through the European Nucleotide Archive study accession number: PRJEB27301.

## References

1 Bozdech, Z. et al. The transcriptome of the intraerythrocytic developmental cycle of Plasmodium falciparum. PLoS Biol 1, E5, doi:10.1371/journal.pbio.0000005 (2003).

2 Le Roch, K. G. et al. Discovery of gene function by expression profiling of the malaria parasite life cycle. Science 301, 1503–1508, doi:10.1126/science.1087025 (2003).

3 Otto, T. D. et al. New insights into the blood-stage transcriptome of Plasmodium falciparum using RNA-Seq. Molecular microbiology 76, 12–24, doi:10.1111/j.1365-2958.2009.07026.x (2010).

4 Sorber, K., Dimon, M. T. & Derisi, J. L. RNA-Seq analysis of splicing in Plasmodium falciparum uncovers new splice junctions, alternative splicing and splicing of antisense transcripts. Nucleic Acids Research 39, 3820–3835, doi:10.1093/nar/gkq1223 (2011).

5 Le Roch, K. G. et al. Global analysis of transcript and protein levels across the Plasmodium falciparum life cycle. Genome Research 14, 2308–2318, doi:10.1101/gr.2523904 (2004).

6 Lu, X. M. et al. Nascent RNA sequencing reveals mechanisms of gene regulation in the human malaria parasite Plasmodium falciparum. Nucleic Acids Research 45, 7825–7840, doi:10.1093/nar/gkx464 (2017).

7 Bunnik, E. M. et al. Polysome profiling reveals translational control of gene expression in the human malaria parasite Plasmodium falciparum. Genome Biology 14, R128, doi:10.1186/gb-2013-14-11-r128 (2013).

8 Caro, F., Ahyong, V., Betegon, M. & DeRisi, J. L. Genome-wide regulatory dynamics of translation in the Plasmodium falciparum asexual blood stages. eLife 3, doi:10.7554/eLife.04106 (2014).

9 Gardner, M. J. et al. Genome sequence of the human malaria parasite Plasmodium falciparum. Nature 419, 498–511, doi:10.1038/nature01097 (2002).

10 Llinas, M., Bozdech, Z., Wong, E. D., Adai, A. T. & DeRisi, J. L. Comparative whole genome transcriptome analysis of three Plasmodium falciparum strains. Nucleic Acids Res 34, 1166–1173, doi:10.1093/nar/gkj517 (2006).

11 Bozdech, Z. et al. The transcriptome of Plasmodium vivax reveals divergence and diversity of transcriptional regulation in malaria parasites. Proc Natl Acad Sci U S A 105, 16290–16295, doi:10.1073/pnas.0807404105 (2008).

12 Lapp, S. A. et al. Plasmodium knowlesi gene expression differs in ex vivo compared to in vitro blood-stage cultures. Malaria Journal 14, 110, doi:10.1186/s12936-015-0612-8 (2015).

13 Oakley, M. S. M. et al. Molecular factors and biochemical pathways induced by febrile temperature in intraerythrocytic Plasmodium falciparum parasites. Infection and Immunity 75, 2012–2025, doi:10.1128/IAI.01236-06 (2007).

14 Natalang, O. et al. Dynamic RNA profiling in Plasmodium falciparum synchronized blood stages exposed to lethal doses of artesunate. BMC Genomics 9, 388, doi:10.1186/1471-2164-9-388 (2008).

15 Hu, G. et al. Transcriptional profiling of growth perturbations of the human malaria parasite Plasmodium falciparum. Nature Biotechnology 28, 91–98, doi:10.1038/nbt.1597 (2010).

16 Ndam, N. T. et al. Plasmodium falciparum transcriptome analysis reveals pregnancy malaria associated gene expression. PLoS ONE 3, doi:10.1371/journal.pone.0001855 (2008).

17 Yamagishi, J. et al. Interactive transcriptome analysis of malaria patients and infecting Plasmodium falciparum. Genome Research 24, 1433–1444, doi:10.1101/gr.158980.113 (2014).

18 Mok, S. et al. Population transcriptomics of human malaria parasites reveals the mechanism of artemisinin resistance. Science 347, 431–435, doi:10.1126/science.1260403 (2015).

19 Langhorne, J. et al. The relevance of non-human primate and rodent malaria models for humans. Malar J 10, 23, doi:10.1186/1475-2875-10-23 (2011).

20 Matz, J. M. & Kooij, T. W. Towards genome-wide experimental genetics in the in vivo malaria model parasite Plasmodium berghei. Pathog Glob Health 109, 46–60, doi:10.1179/2047773215Y.0000000006 (2015).

21 Stephens, R., Culleton, R. L. & Lamb, T. J. The contribution of Plasmodium chabaudi to our understanding of malaria. Trends Parasitol 28, 73–82, doi:10.1016/j.pt.2011.10.006 (2012).

22 Hall, N. et al. A comprehensive survey of the Plasmodium life cycle by genomic, transcriptomic, and proteomic analyses. Science (New York, N.Y.) 307, 82–86, doi:10.1126/science.1103717 (2005).

23 Otto, T. D. et al. A comprehensive evaluation of rodent malaria parasite genomes and gene expression. BMC biology 12, 86, doi:10.1186/PREACCEPT-1233682211145405 (2014).

24 Hoo, R. et al. Integrated analysis of the Plasmodium species transcriptome. EBioMedicine 7, 255–266, doi:10.1016/j.ebiom.2016.04.011 (2016).

25 Mair, G. R. Regulation of Sexual Development of Plasmodium by Translational Repression. Science 313, 667–669, doi:10.1126/science.1125129 (2006).

26 Mair, G. R. et al. Universal features of post-transcriptional gene regulation are critical for Plasmodium zygote development. PLoS Pathogens 6, doi:10.1371/journal.ppat.1000767 (2010).

27 Silvie, O., Briquet, S., Müller, K., Manzoni, G. & Matuschewski, K. Post-transcriptional silencing of UIS4 in Plasmodium berghei sporozoites is important for host switch. Molecular Microbiology 91, 1200–1213, doi:10.1111/mmi.12528 (2014).

28 Tarun, A. S. et al. A combined transcriptome and proteome survey of malaria parasite liver stages. Proc Natl Acad Sci U S A 105, 305–310, doi:10.1073/pnas.0710780104 (2008).

29 Lawton, J. et al. Characterization and gene expression analysis of the cir multi gene family of Plasmodium chabaudi chabaudi (AS). BMC Genomics 13, 125, doi:10.1186/1471-2164-13-125 (2012).

30 Brugat, T. et al. Antibody-independent mechanisms regulate the establishment of chronic Plasmodium infection. Nat Microbiol 2, 16276, doi:10.1038/nmicrobiol.2016.276 (2017).

31 Lin, J.-w. et al. Signatures of malaria-associated pathology revealed by high resolution whole-blood transcriptomics in a rodent model of malaria. Scientific Reports 7, 41722, doi:10.1038/srep41722 (2017).

32 Spence, P. J. et al. Vector transmission regulates immune control of Plasmodium virulence. Nature 498, 228–231, doi:10.1038/nature12231 (2013).

33 Miller, Jessica L., Sack, Brandon K., Baldwin, M., Vaughan, Ashley M. & Kappe, Stefan H. I. Interferon-Mediated Innate Immune Responses against Malaria Parasite Liver Stages. Cell Reports 7, 436–447, doi:10.1016/j.celrep.2014.03.018 (2014).

34 Lindner, S. E. et al. Perturbations of PlasmodiumPuf2 expression and RNA-seq of Puf2-deficient sporozoites reveal a critical role in maintaining RNA homeostasis and parasite transmissibility. Cellular Microbiology 15, 1266–1283, doi:10.1111/cmi.12116 (2013).

35 Roques, M. et al. Plasmodium P-Type Cyclin CYC3 Modulates Endomitotic Growth during Oocyst Development in Mosquitoes. PLoS pathogens 11, e1005273, doi:10.1371/journal.ppat.1005273 (2015).

36 Guttery, D. S., Roques, M., Holder, A. A. & Tewari, R. Commit and Transmit: Molecular Players in Plasmodium Sexual Development and Zygote Differentiation. Trends Parasitol 31, 676–685, doi:10.1016/j.pt.2015.08.002 (2015).

37 Modrzynska, K. et al. A Knockout Screen of ApiAP2 Genes Reveals Networks of Interacting Transcriptional Regulators Controlling the Plasmodium Life Cycle. Cell Host and Microbe 21, 11–22, doi:10.1016/j.chom.2016.12.003 (2017).

38 Shaw, P. J. et al. Plasmodium parasites mount an arrest response to dihydroartemisinin, as revealed by whole transcriptome shotgun sequencing (RNA-seq) and microarray study. BMC Genomics 16, 830, doi:10.1186/s12864-015-2040-0 (2015).

39 Parasuraman, S., Raveendran, R. & Kesavan, R. Blood sample collection in small laboratory animals. Journal of pharmacology & pharmacotherapeutics 1, 87–93, doi:10.4103/0976-500X.72350 (2010).

40 Venkatesan, M. et al. Using CF11 cellulose columns to inexpensively and effectively remove human DNA from Plasmodium falciparum-infected whole blood samples. Malaria Journal 11, 41, doi:10.1186/1475-2875-11-41 (2012).

41 Chomczynski, P. A reagent for the single-step simultaneous isolation of RNA, DNA and proteins from cell and tissue samples. BioTechniques 15, 532–537, doi:10.1038/news.2010.498 (1993).

42 Chomczynski, P. & Sacchi, N. Single-step method of RNA isolation by acid guanidinium thiocyanate-phenol-chloroform extraction. Analytical Biochemistry 162, 156–159, doi:10.1016/0003-2697(87)90021-2 (1987).

43 Zhu, L. et al. New insights into the Plasmodium vivax transcriptome using RNA-Seq. Scientific Reports 6, 20498, doi:10.1038/srep20498 (2016).

44 Siegel, T. N. et al. Strand-specific RNA-Seq reveals widespread and developmentally regulated transcription of natural antisense transcripts in Plasmodium falciparum. BMC genomics 15, 150, doi:10.1186/1471-2164-15-150 (2014).

45 Bateman, K. P. et al. Reduction of animal usage by serial bleeding of mice for pharmacokinetic studies: Application of robotic sample preparation and fast liquid chromatography-mass spectrometry. Journal of Chromatography B: Biomedical Sciences and Applications 754, 245–251, doi:10.1016/S0378-4347(00)00612-5 (2001).

46 Peng, S. X., Rockafellow, B. A., Skedzielewski, T. M., Huebert, N. D. & Hageman, W. Improved pharmacokinetic and bioavailability support of drug discovery using serial blood sampling in mice. Journal of Pharmaceutical Sciences 98, 1877–1884, doi:10.1002/jps.21533 (2009).

47 Rahavendran, S. V. et al. Discovery pharmacokinetic studies in mice using serial microsampling, dried blood spots and microbore LC-MS/MS. Bioanalysis 4, 1077–1095, doi:10.4155/bio.12.85 (2012).

48 Watanabe, A. et al. Using improved serial blood sampling method of mice to study pharmacokinetics and drug-drug interaction. Journal of Pharmaceutical Sciences 104, 955–961, doi:10.1002/jps.24236 (2015).

49 Gautret, P., Deharo, E., Chabaud, A. G., Ginsburg, H. & Landau, I.Plasmodium vinckei vinckei P. v. lentum and P. yoelii yoelii: chronobiology of the asexual cycle in the blood. Parasite 1, 235–239, doi:10.1051/parasite/1994013235 (1994).

50 Ramaprasad, A. Phenomics, Genomics and Genetics in Plasmodium vinckei PhD thesis, King Abdullah University of Science and Technology, (2017).

51 Noedl, H. et al. Evidence of artemisinin-resistant malaria in western Cambodia. N Engl J Med 359, 2619–2620, doi:10.1056/NEJMc0805011 (2008).

52 Patel, N. J. et al. Evaluation and Optimization of Blood Micro-Sampling Methods: Serial Sampling in a Cross-Over Design from an Individual Mouse. J Pharm Pharm Sci 19, 496–510, doi:10.18433/J3NK60 (2016).

53 Korfmacher, W. et al. Capillary microsampling of whole blood for mouse PK studies: an easy route to serial blood sampling. Bioanalysis 7, 449–461, doi:10.4155/bio.14.275 (2015).

54 Kim, D. et al. TopHat2: accurate alignment of transcriptomes in the presence of insertions, deletions and gene fusions. Genome Biol 14, R36, doi:10.1186/gb-2013-14-4-r36 (2013).

55 Trapnell, C. et al. Differential analysis of gene regulation at transcript resolution with RNA-seq. Nat Biotechnol 31, 46–53, doi:10.1038/nbt.2450 (2013).

56 Wang, L., Wang, S. & Li, W. RSeQC: quality control of RNA-seq experiments. Bioinformatics 28, 2184–2185, doi:10.1093/bioinformatics/bts356 (2012).

57 Becker, R. A., Chambers, J. M. & Wilks, A. R. The New S Language. Pacific Grove Ca Wadsworth Brooks 1988 1, 702 (1988).

58 Wei, T. & Simko, V. corrplot: Visualization of a Correlation Matrix. (2016).

59 Yang, R. & Su, Z. Analyzing circadian expression data by harmonic regression based on autoregressive spectral estimation. Bioinformatics 26, i168–174, doi:10.1093/bioinformatics/btq189 (2010).

60 Warnes, G. R. et al. gplots: Various R Programming Tools for Plotting Data. (2016).

61 Wickham, H. ggplot2: Elegant Graphics for Data Analysis. (2009).

62 Liao, Y., Smyth, G. K. & Shi, W. FeatureCounts: An efficient general purpose program for assigning sequence reads to genomic features. Bioinformatics 30, 923–930, doi:10.1093/bioinformatics/btt656 (2014).

63 Robinson, D. G. & Storey, J. D. SubSeq: Determining appropriate sequencing depth through efficient read subsampling. Bioinformatics 30, 3424–3426, doi:10.1093/bioinformatics/btu552 (2014).

64 Love, M. I., Anders, S. & Huber, W. Differential analysis of count data - the DESeq2 package. Genome Biology 15, 550, doi:110.1186/s13059-014-0550-8 (2014).

65 Robinson, M. D., McCarthy, D. J. & Smyth, G. K. edgeR: a Bioconductor package for differential expression analysis of digital gene expression data. Bioinformatics (Oxford, England) 26, 139–140, doi:10.1093/bioinformatics/btp616 (2010).

66 Goodwin, S., McPherson, J. D. & McCombie, W. R. Coming of age: ten years of next-generation sequencing technologies. Nat Rev Genet 17, 333–351, doi:10.1038/nrg.2016.49 (2016).

